# Dual role of RBM3 in PEDV infection: stabilization of viral RNA and upregulation of the entry factor HSPA5

**DOI:** 10.64898/2026.11.02.763835

**Authors:** Siyu Li, Yan Zeng, Jinze Li, Yuchang Liu, Huili Liu, Xinyu Yang, Wenzhen Qin, Tao Li, Zongyan Chen, Jingyi Liu, Guangzhi Tong, Xin Cao, Ning Kong, Tongling Shan

## Abstract

RBM3, a member of the cold shock protein family, functions as an RNA chaperone that stabilizes target transcripts by binding to their untranslated regions (UTRs). Porcine epidemic diarrhea virus (PEDV) is an enteric coronavirus that poses a serious threat to the swine industry, and its replication heavily depends on its genomic and subgenomic RNAs, making these RNAs potential targets for host RNA binding proteins. In this study, we show that RBM3 promotes PEDV replication through a dual mechanism. On one hand, RBM3 exhibits a binding preference for the viral 3’UTR, which prolongs the half-life of viral RNA and thereby promotes PEDV proliferation. RBM3 also interacts with the PEDV nucleocapsid (N) protein and replication-associated double-stranded RNA (dsRNA) signals, suggesting that RBM3 may be associated with viral replication complexes, though its contribution to their functional integrity remains to be determined. On the other hand, RBM3 increases HSPA5 promoter activity and HSPA5 mRNA levels, linking RBM3 to HSPA5, a host factor known to facilitate PEDV attachment and internalization. These two mechanisms together provide a mechanistic basis for efficient PEDV infection. Our findings identify RBM3 as a positive host regulator of PEDV and establish, for the first time, a functional link between RBM3 and HSPA5, offering new insights into coronavirus host interactions and suggesting RBM3 as a potential target for antiviral intervention.

## 1. Introduction

Porcine epidemic diarrhea virus (PEDV) is an enteric alphacoronavirus that poses a serious threat to the swine industry [1]. Its genome is a single-stranded positive-sense RNA of approximately 28 kb, containing a 5’untranslated region (UTR), two overlapping replicase open reading frames (ORF1a and ORF1b), genes encoding structural proteins (S, E, M, and N), and a 3’UTR [1]. Among these, the nucleocapsid (N) protein is the most abundant viral structural protein and plays a multifaceted role beyond genome encapsidation [2]. For instance, the PEDV N protein interacts with host proteins DDX5 and SND1, as well as with viral non-structural proteins nsp9 and nsp12, to facilitate the assembly of the replication-transcription complex (RTC) and promote viral RNA synthesis [3]. Additionally, the 3’UTR, particularly its bulged stem-loop structure, serves as a cis-acting element required for N-protein-mediated enhancement of viral RNA translation [4], highlighting this region as a potential target for host factor recognition. Although these findings underscore the importance of viral RNA elements and associated proteins in PEDV replication, how host RNA-binding proteins (RBPs) recognize and regulate PEDV RNA components remains largely unexplored.

RNA-binding proteins (RBPs) are key regulators of post-transcriptional gene expression, influencing RNA splicing, transport, stability, and translation by recognizing specific sequences or structural elements in target transcripts [5, 6]. RNA-binding motif protein 3 (RBM3), a member of the cold-shock protein family, contains an N-terminal RNA recognition motif (RRM) and a C-terminal glycine-rich domain [7]. Under stress conditions such as hypothermia and hypoxia, RBM3 is upregulated and functions as an RNA chaperone that stabilizes target mRNAs by binding to their 3’UTRs [8]. For example, RBM3 binds to the 3’UTR of Yap1 mRNA to enhance its stability, thereby protecting neurodevelopment from cold-induced damage [9]. Despite its well-characterized roles in cellular RNA metabolism, whether RBM3 participates in viral RNA regulation (particularly during coronavirus infection) remains unknown. Interestingly, a recent transcriptomic study reported that RBM3 and HSPA5 exhibit similar expression patterns in the mouse brain following sleep deprivation [10], hinting at a potential regulatory link between these two genes. HSPA5 (also known as GRP78/BiP) is an endoplasmic reticulum-resident chaperone involved in protein folding, and has also been identified as a cell-surface attachment and entry factor for several coronaviruses, including SARS-CoV-2 [11] and PEDV [12]. In PEDV infection, HSPA5 specifically binds to the spike (S) protein and promotes viral attachment and internalization [12]. These observations prompted us to investigate whether RBM3, beyond its direct effects on viral RNA, might also influence PEDV infection through regulation of HSPA5.

In this study, we investigated the role of RBM3 in the PEDV life cycle. We found that RBM3 promotes PEDV infection through two distinct routes. Directly, RBM3 binds preferentially to the viral 3’UTR, which prolongs viral RNA half-life and enhances replication. Interestingly, it also associates with the N protein and dsRNA. Indirectly, RBM3 increases the promoter activity and mRNA levels of HSPA5, a host protein known to facilitate viral entry. These findings identify RBM3 as a positive host regulator of PEDV and reveal an unexpected functional connection between RBM3 and HSPA5, providing new insights into how a single host RBP can coordinate both viral RNA metabolism and host protein regulation during coronavirus infection.

## 2. Materials and methods

### 2.1 Cell culture and viruses

LLC-PK1 cells were cultured in MEM, while HEK293T and HeLa cells were cultured in DMEM, both media supplemented with 10% heat-inactivated FBS, at 37°C in a humidified incubator with 5% CO . The PEDV strain used was the laboratory-isolated PEDV variant JS-2013 (MH910099).

### 2.2 Antibodies and reagents

The following primary antibodies were used in this study: anti-β-actin (Proteintech, 66009-1-Ig), anti-Flag (Sigma-Aldrich, F1804), anti-dsRNA (Sigma-Aldrich, MABE113), and an in-house custom antibody against the PEDV N protein. Horseradish peroxidase (HRP)-conjugated anti-mouse (Proteintech, SA00001-1) and anti-rabbit (Proteintech, SA00001-2) were used as secondary antibodies. The inhibitors were Remdesivir (MCE, GS-5734) and Actinomycin D (MCE, HY-17559).

### 2.3 Plasmids and transfection

RBM3 (XM_005657795.4) and HSPA5 (NM_001444629.1) with a C-terminal Flag tag were cloned into the pXJ41 vector using the ClonExpress II One Step Cloning Kit (Vazyme Biotech, C112-01). PEDV N and NSP12 plasmids with a C-terminal HA tag were stored in our laboratory. For cellular delivery, the plasmids were transfected into cells at 80–90% confluence using Lipofectamine 3000 transfection reagent (Invitrogen, L3000015), and siRNAs were transfected into cells at 60-70% confluence using Lipofectamine RNAiMAX (Invitrogen, 13778030).

### 2.4 RNA immunoprecipitation (RIP)

LLC-PK1 cells overexpressing Flag-RBM3 were infected with PEDV 24 h after transfection. Samples were collected at 4 h post-infection and temporarily stored on ice. Magnetic beads were pre-coupled with IgG or Flag antibodies overnight, then rinsed with NT2 buffer to remove excess antibodies. Equal amounts of sample were added to tubes containing IgG or Flag antibody-conjugated beads, along with RNA capture buffer, and incubated at 4°C for 4–6 h. After incubation, the beads were washed with NT2 buffer to remove unbound materials. RNA was extracted using the Trizol method, immediately reverse transcribed into cDNA, and subsequently analyzed by qPCR to assess RNA pull-down efficiency.

### 2.5 RNA pull-down

For the generation of in vitro transcription templates, primer pairs were designed with the forward primer incorporating a T7 promoter sequence. Following amplification, the PCR products were purified and subsequently employed as templates for transcription. In vitro synthesis of RNA was conducted utilizing the EasyCap T7 Co-Transcription Kit (Vazyme Biotech, DD4203), which includes a CAG trimer. The resultant RNA transcripts were purified using VAHTS RNA Clean Beads (Vazyme Biotech, N412). Finally, the purified RNA underwent 3’-end desthiobiotinylation using the Pierce RNA 3’End Desthiobiotinylation Kit (Thermo Fisher Scientific, 20163) prior to storage at -80°C.

HEK293T cells were cultured in T25 flasks until they reached approximately 90% confluence. The cells were then transfected with a plasmid encoding Flag-tagged RBM3. At 24 h post-transfection, cells were lysed on ice for 30 min using Pierce IP Lysis Buffer (Thermo Fisher Scientific, 87787) supplemented with a protease inhibitor cocktail, followed by cell scraping. The lysate was centrifuged at 12,000 rpm for 5 min at 4°C, and the resulting supernatant was collected and stored at −80°C for subsequent analysis. To capture RNA-binding proteins (RBPs), the labeled RNA was incubated with the prepared lysate using the Pierce Magnetic RNA-Protein Pull-Down Kit (Thermo Fisher Scientific, 20164) according to the manufacturer’s instructions. Finally, the eluted RBPs were mixed with loading buffer, heated for denaturation, separated from the magnetic beads, and subjected to Western blotting analysis.

### 2.6 Co-immunoprecipitation (Co-IP)

HEK293T cells were transfected with the indicated plasmids. At 24 h post-transfection, protein samples were harvested and stored at –80 °C. Blank magnetic beads (Thermo Fisher, 88804) were washed 2–3 times with PBST, followed by incubation with the primary antibody at room temperature for 50 min. After incubation, the beads were washed 4–5 times with PBST. The frozen protein samples were thawed on ice and then incubated with the antibody-bound beads at room temperature for 90 min. Subsequently, the beads were washed 5–6 times with PBST, and the residual liquid was removed. Glycine elution buffer was added, and the mixture was vortexed thoroughly. After a 1-minute incubation, protein loading buffer was added to denature the samples. If the protein of interest is an RNA-binding protein, prior to elution, the magnetic beads should be equally divided into two aliquots. One aliquot is directly eluted, while the other is incubated with 1 μL of RNase A for an additional 30 minutes before proceeding with elution. The resulting samples were then analyzed by Western blotting.

### 2.7 Confocal Immunofluorescence assay

For immunofluorescence staining, cells were first fixed with 4% paraformaldehyde (Sigma-Aldrich, P6148) for 15 min at room temperature and then permeabilized with 0.1% Triton X-100 (Sigma-Aldrich, T9284) for 10 min. After blocking with 5% bovine serum albumin for 1 hour at 37°C, the cells were incubated with primary antibodies, followed by fluorophore-conjugated secondary antibodies, according to previously established protocols. Nuclei were counterstained with 4’,6-diamidino-2-phenylindole (DAPI; Beyotime Biotechnology, C1002) for 5 min. Subsequently, the fluorescence was observed using a Zeiss Laser Scanning Confocal Microscope.

### 2.8 RNA stability assay

LLC-PK1 cells overexpressing Flag-RBM3 or the empty vector were infected with PEDV at 24 hours post-transfection. At 8 hours post-infection, cells were treated with 20 μM Remdesivir, and samples were collected at various time points to identify the time point (T0) at which viral RNA synthesis plateaued or ceased, as determined by RT-qPCR. After identifying T0, the experiment was repeated. Starting from T0, samples were collected every two hours and stored at −80°C for subsequent RT-qPCR analysis. RNA half-life calculation: RNA degradation was assumed to follow a first-order exponential decay model: *Y* = *A* · *e*^(-*kt*), where *y* is the normalized relative RNA amount, *t* is time (h), and *k* is the degradation rate constant. Taking the natural logarithm of both sides gives the linear form *ln* y = *ln A* - *kt*. Thus, a linear regression of ln *y* versus *t* was performed, the slope *b* yielding *K* = -*b*, and the half-life was calculated as *t*_ (1/2) = *ln* 2/*k*.

### 2.9 Dual-luciferase reporter assay

HEK293T cells were co-transfected with the firefly luciferase reporter plasmid and the Renilla luciferase internal control plasmid. At 24 h post-transfection, cells were lysed using the lysis buffer provided in the Dual Luciferase Reporter Assay Kit (Vazyme, Catalog No. DL101-01). The lysates were centrifuged at room temperature for 2 min, and the supernatants were collected. A 100 µL aliquot of luciferase substrate (equilibrated to room temperature) was dispensed into each well of a microplate. Then, 20 µL of the supernatant was carefully added to the well, mixed rapidly, and firefly luciferase activity was immediately measured using a microplate reader. Subsequently, 100 µL of freshly prepared Renilla substrate working solution was added to the same reaction mixture, mixed rapidly, and Renilla luciferase activity was measured immediately. The final results were calculated as the ratio of firefly luciferase activity to Renilla luciferase activity.

### 2.10 Western blotting assay

Protein lysates, prepared using RIPA buffer (Thermo Fisher Scientific, 89901) with protease inhibitors, were mixed with loading buffer and denatured at 100°C for 10 min. After SDS-PAGE separation, proteins were transferred to NC membranes (GE Healthcare, 10600001) and immunoblotted with indicated primary and secondary antibodies. Signals were visualized using an ECL kit (LumiQ, Share-bio, SB-WB012).

### 2.11 RNA isolation and RT-qPCR

Total RNA was isolated with Trizol reagent. In brief, cell monolayers were disrupted in 1 mL of Trizol for 5 min. The homogenate was mixed with 200 µL of chloroform by inversion, incubated for 3 min, and then centrifuged at 4°C for 15 min. The clear upper phase was transferred to a fresh tube, and RNA was precipitated by adding 500 µL of isopropanol. Following centrifugation, the RNA pellet was washed twice with 75% ethanol, air-dried, and resuspended in RNase-free water. Reverse transcription was conducted with the HiScript III RT SuperMix for qPCR (Vazyme Biotech, R323–01). The resulting cDNA was then subjected to quantitative PCR analysis on a Roche LightCycler 480 instrument using SYBR Premix Ex Taq (Vazyme Biotech, q711–03), following the manufacturer’s instructions.

### 2.12 CCK-8

After the cells had adhered, inhibitors were added according to the experimental design, followed by incubation for 12 h. CCK-8 reagent (Share Bio, SB-CCK8) was then added to each well at a volume equivalent to 10% of the culture medium volume. The cells were incubated for an additional 2 h in a humidified cell culture incubator, and absorbance was measured at 450 nm using a microplate reader. Relative cell viability was calculated using the following equation:

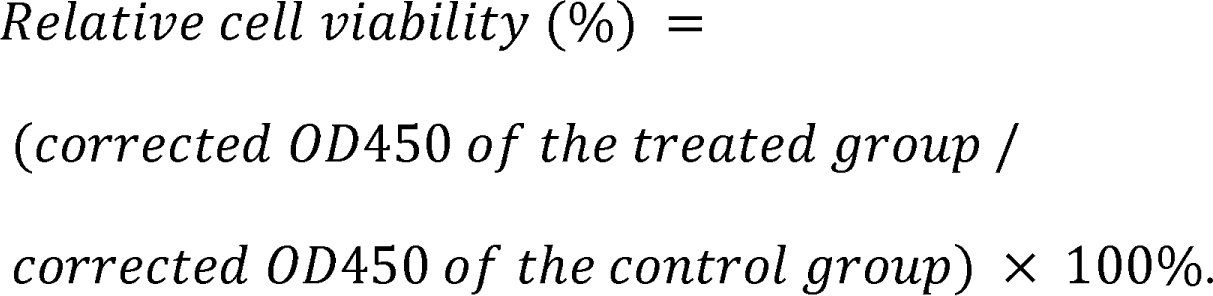

### 2.13 Statistical analysis

We used GraphPad Prism 5.0 software for all statistical analyses, with results presented as the mean ± SD. Differences between groups were evaluated using two-way ANOVA or unpaired t-test. Statistical significance is indicated in the figures as follows: *P ≤ 0.05, **P ≤ 0.01, ***P ≤ 0.001; ns denotes not significant (P > 0.05).

## 3. Results

### 3.1 RBM3 promotes PEDV proliferation

Based on our laboratory’s previous proteomics screening results, we identified a set of candidate proteins potentially involved in the PEDV proliferation process (Ref. [3], Fig.1B). Among these candidates, RBM3 was selected for further investigation. To investigate the effect of RBM3 on PEDV replication, we overexpressed Flag-RBM3 in LLC-PK1 cells, infected them with PEDV 24 h later, and collected cell samples at different time points post-infection. Analysis by Western blotting (Fig. 1A) and RT-qPCR (Fig. 1B) confirmed the promoting effect of RBM3 on PEDV proliferation. Subsequently, we transfected increasing doses of RBM3 before PEDV infection, and the results showed that both PEDV N protein expression (Fig. 1C) and viral RNA levels (Fig. 1D) increased in a dose-dependent manner following RBM3 transfection. Additionally, we constructed a cell line stably overexpressing RBM3. Functional validation using this cell line demonstrated that RBM3 overexpression significantly promotes PEDV proliferation, as evidenced by Western blotting (Fig. 1E) and RT-qPCR (Fig. 1F) results. In contrast, RBM3 knockdown significantly inhibited PEDV proliferation. The knockdown efficiency of si-RBM3 was confirmed by RT-qPCR (Fig. 1G). Accordingly, RBM3 depletion reduced PEDV N protein levels, as determined by Western blotting (Fig. 1H), and viral RNA levels, as determined by RT-qPCR (Fig. 1I).

**Figure 1.**
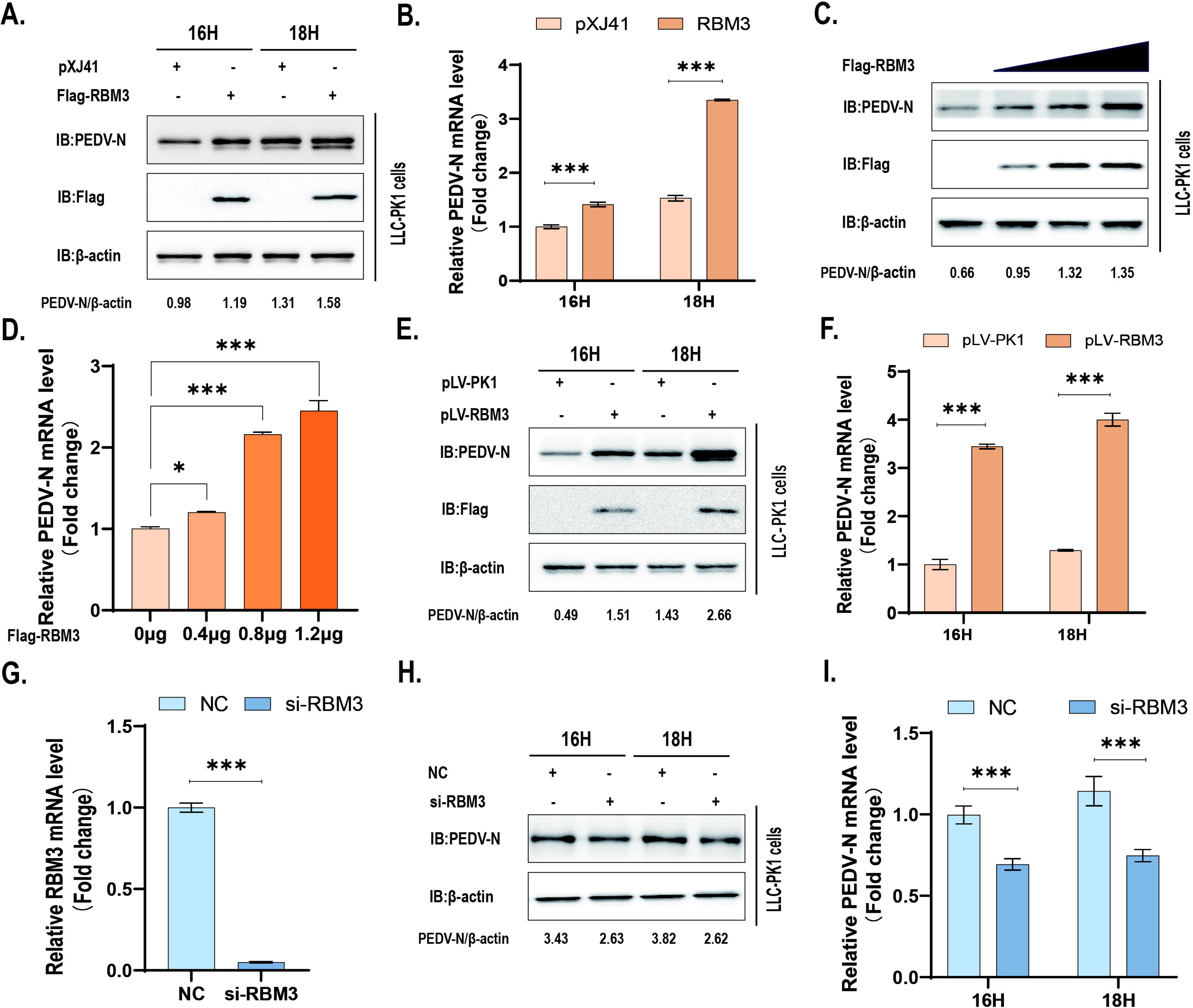
RBM3 promotes PEDV proliferation. (A, B) Western blotting and RT qPCR analysis of cells overexpressing Flag RBM3 versus empty vector control at indicated hours post infection (hpi) with PEDV. P-values determined by two-way ANOVA. (C, D) Western blotting and RT qPCR analysis of cells transfected with increasing doses of Flag RBM3 plasmid (0, 0.4, 0.8, 1.2 μg) and infected with PEDV. P-values determined by two-way ANOVA. (E, F) Western blotting and RT qPCR analysis of stable RBM3 overexpressing cell line (RBM3 OE) versus empty vector control at indicated hpi with PEDV. P-values determined by two-way ANOVA. (G) Assessment of the interference efficiency of si-RBM3. P-values determined by unpaired t-test. (H, I) Western blotting and RT qPCR analysis of cells transfected with si RBM3 versus NC and infected with PEDV at indicated hpi. All data represent three independent biological replicates (n=3). Error bars represent mean ± SD. *p < 0.05, **p < 0.01, ***p < 0.001.

### 3.2 RBM3 preferentially binds the 3’UTR of PEDV RNA and prolongs its half-life

Given the RNA-binding properties of RBM3, we infected Flag-RBM3-overexpressing cells with PEDV and performed RIP-qPCR assays. The results revealed that RBM3 binds to PEDV RNA (Fig. 2A). To further map the binding site of RBM3 on the viral RNA, we cloned the non-coding regions of the PEDV genome (5’UTR and 3’UTR) into plasmids. These constructs were separately transfected into Flag-RBM3-overexpressing cells, and after 24 h, RIP-qPCR analysis demonstrated that RBM3 is enriched at the 3’UTR of PEDV RNA (Fig. 2B). To validate these findings, 5’UTR and 3’UTR RNAs were transcribed in vitro and biotinylated. A biotin-labeled scrambled RNA was used as a negative control. Flag RBM3 protein was pulled down by RNA pull down assays and subsequently detected by Western blotting. The results showed that the 3’UTR efficiently pulled down Flag RBM3 protein (Fig. 2C). To verify the functional relevance of this binding preference and to establish the conditions for subsequent viral RNA half-life analysis, we performed a series of functional validation and preliminary experiments. First, we examined whether the RBM3-mediated increase in viral RNA abundance depends on the 3’UTR sequence: Cells with either RBM3 overexpression or knockdown were co-transfected with equal amounts of a plasmid containing the viral 3’UTR sequence. RT-qPCR results showed that, compared to the control group, RBM3 overexpression significantly increased the abundance of 3’UTR RNA, whereas RBM3 knockdown markedly reduced it. However, when the PEDV N plasmid was used instead, this effect was not observed (Fig. S1A). These results suggest that the RBM3-mediated effect is not a general consequence of nonspecific viral RNA stabilization, but is associated with the presence of the 3’UTR sequence. Next, we assessed cell viability after Remdesivir treatment by CCK-8 assay (Fig. S1B) and determined the time point at which viral RNA synthesis was completely inhibited (T0), thereby optimizing the conditions for the subsequent RNA half-life analysis: We infected the Flag-RBM3 overexpressing cell line with PEDV and then added the viral RNA-dependent RNA polymerase (RdRp) inhibitor Remdesivir (20 µM) at 8 h post-infection to halt ongoing RNA synthesis. Samples were collected at different time points after drug treatment and analyzed by RT-qPCR. The results showed that viral RNA synthesis reached a plateau between 6 and 9 h after Remdesivir treatment (Fig. S1C). To ensure complete inhibition of RNA synthesis, 9 h was selected as T0 in subsequent experiments. After confirming the functional relevance of this binding preference and defining T0 as the time point of complete inhibition of viral RNA synthesis, we performed the viral RNA half-life assay. Using this optimized condition, we repeated the infection and Remdesivir treatment and collected samples at 0, 2, 4 and 6 h after T0. RT-qPCR analysis of these samples was used to generate the viral RNA decay curve. The calculated degradation rate constant and half-life in the experimental group were significantly lower and significantly longer, respectively, than those in the control group (Fig. 2D). Having established that RBM3 overexpression prolongs viral RNA half-life, we next examined its effect on viral RNA accumulation. Cells with RBM3 overexpression or knockdown were infected with PEDV and samples were collected at 1, 2, 3 and 4 h post-infection. We found that RBM3 overexpression markedly increased viral RNA levels (Fig. 2E), whereas RBM3 knockdown led to a corresponding decrease (Fig. 2F). Collectively, these results indicate that RBM3 mediated prolongation of viral RNA half life leads to a significantly higher accumulation of PEDV RNA at early stages of infection compared with that in the corresponding controls.

**Figure 2.**
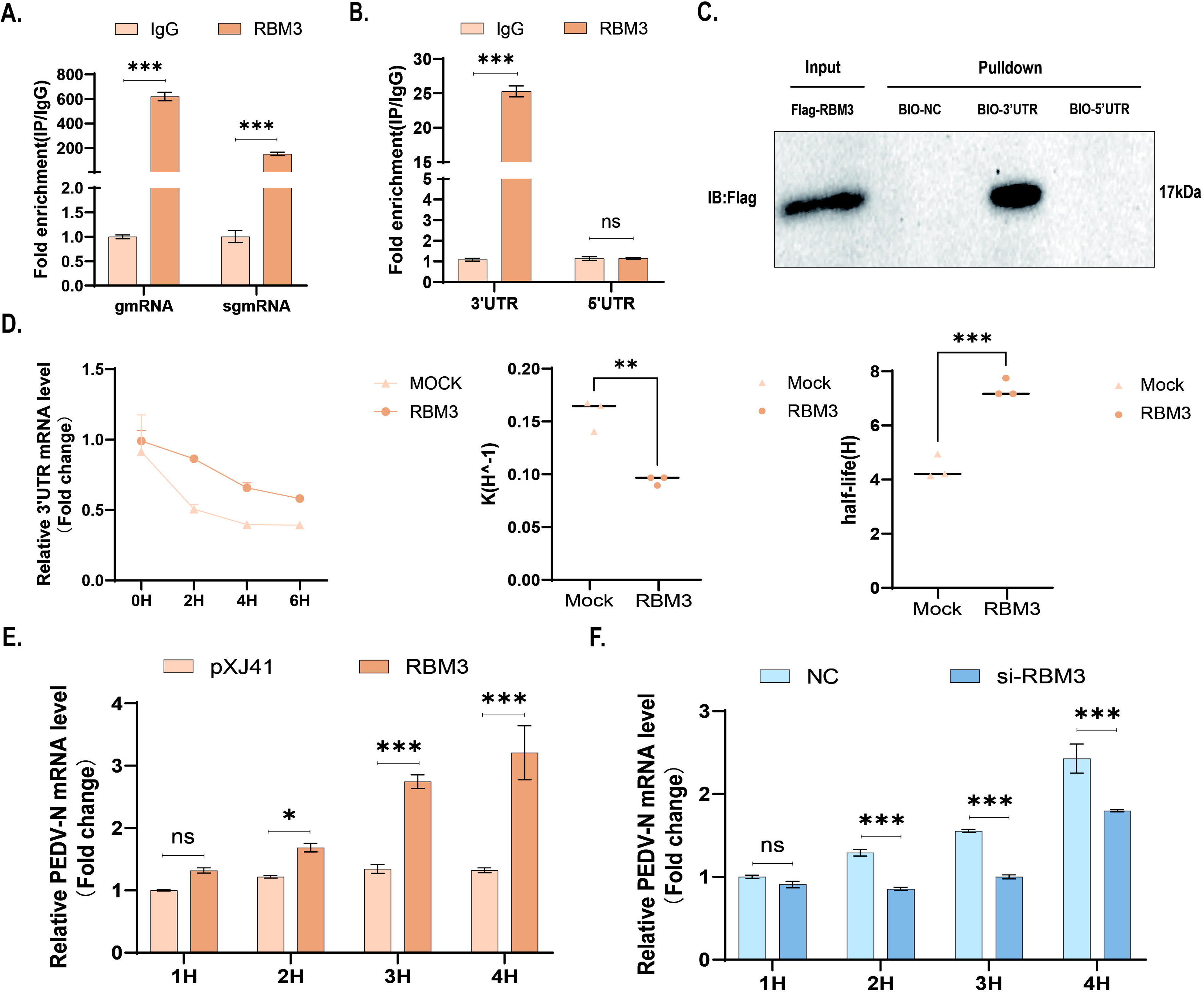
RBM3 preferentially binds the 3’UTR of PEDV RNA and prolongs its half-life. (A) RIP qPCR analysis of RBM3 binding to PEDV RNA in Flag RBM3 overexpressing cells upon PEDV infection. P-values determined by two-way ANOVA. (B) In Flag RBM3 overexpressing cells transfected with plasmids harboring the PEDV 5’UTR or 3’UTR fragments, RIP qPCR was performed to assess RBM3 occupancy on each fragment, revealing preferential enrichment at the 3’UTR. P-values determined by two-way ANOVA. (C) Biotin labeled 5’UTR and 3’UTR RNAs, along with a scrambled control RNA provided in the kit, were generated by in vitro transcription and used in RNA pull down assays to pull down RBM3 protein, Western blotting revealed that only the 3’UTR RNA yielded successful retrieval.(D) Following PEDV infection in RBM3 overexpressing cells and vector control cells, Remdesivir (20 µM) was added to block de novo RNA synthesis, samples were harvested every two hours after the T0 time point for RT qPCR analysis. Viral RNA decay curves were generated. The was significantly lower and the half life of viral RNA was significantly prolonged in RBM3 overexpressing cells compared with control cells. P-values determined by unpaired t-test. (E, F) Cells transfected with Flag RBM3, si RBM3, or vector control were infected with PEDV, and viral RNA accumulation was monitored over 1–4 h post infection. RT qPCR results showed that viral RNA levels were significantly higher in RBM3 overexpressing cells than in the control group, whereas they were significantly lower upon RBM3 knockdown. P-values determined by two-way ANOVA. All data represent three independent biological replicates (n=3). Error bars represent mean ± SD. *p < 0.05, **p < 0.01, ***p < 0.001.

### 3.3 RBM3 interacts with the PEDV N protein and dsRNA

To explore whether RBM3 interacts with other PEDV components beyond previously characterized targets, Co-IP assays were independently performed in LLC-PK1 and HEK293T cells after each cell line was co-transfected with Flag-tagged RBM3 and HA-tagged PEDV N constructs. Western blotting analysis demonstrated that RBM3 was able to pull down PEDV N protein, confirming their physical association (Fig. 3A, B). To further validate this interaction in a cellular context, Flag-RBM3 and HA-PEDV N were separately co-transfected into HeLa and LLC-PK1 cells, and confocal laser scanning microscopy was subsequently employed. Imaging results revealed distinct co-localization signals between RBM3 and PEDV N, providing independent evidence supporting their in-cell interaction (Fig. 3C, D). Having established the interaction between RBM3 and PEDV N protein, we next investigated whether RBM3 could also interact with viral dsRNA, a molecular hallmark of active viral replication. Therefore, we overexpressed Flag-RBM3 in LLC-PK1 cells and infected the cells with PEDV 24 h later. At 4 h post-infection, confocal microscopy revealed the co-localization of RBM3 with dsRNA (Fig. 3E). Although the interactions of RBM3 with PEDV N protein and dsRNA suggested a potential role in the viral replication cycle, Co-IP analysis did not reveal any association between RBM3 and NSP12, a key subunit of the RTC (Fig. 3F).

**Figure 3.**
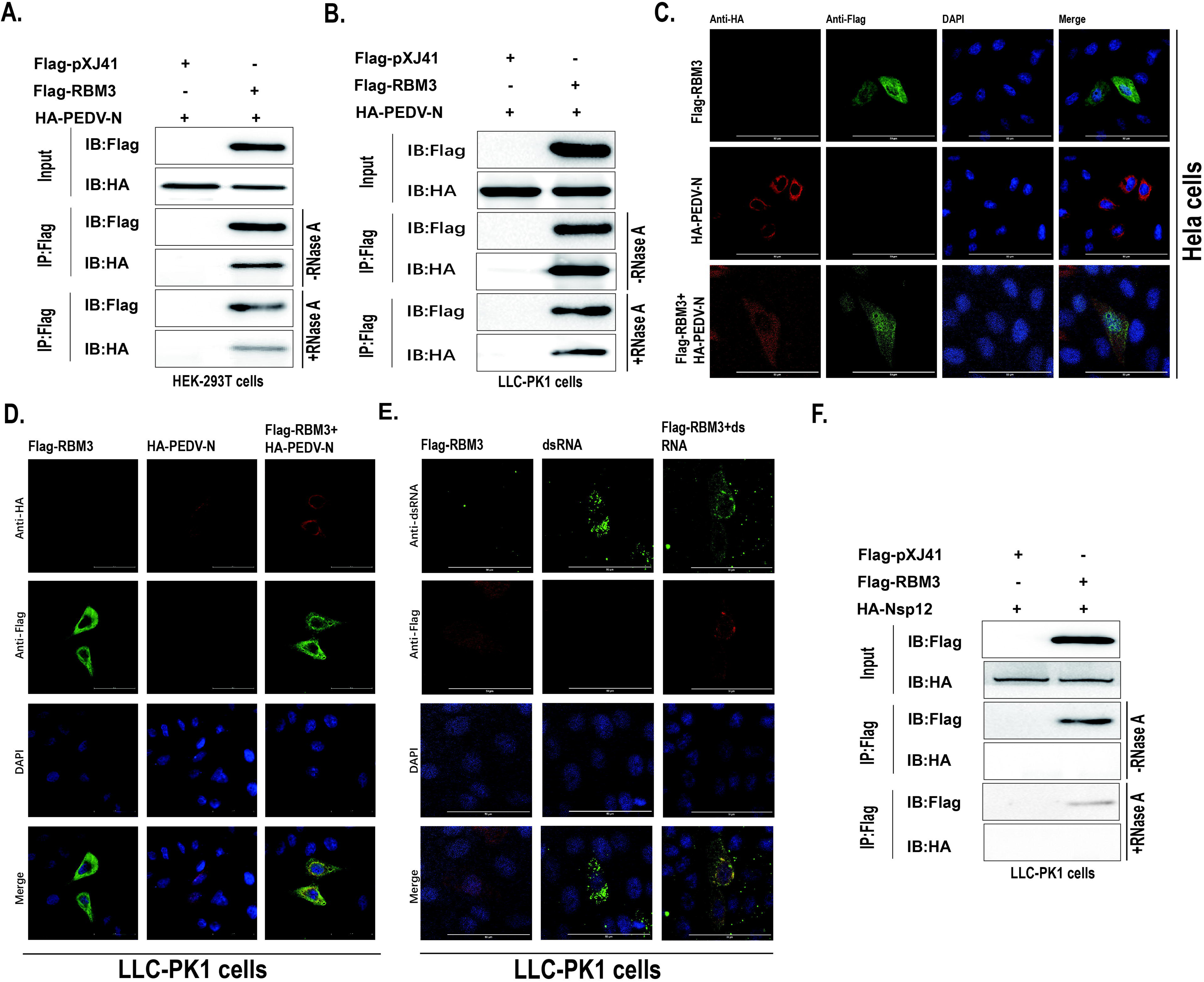
RBM3 interacts with the PEDV N protein and dsRNA. (A, B) Flag RBM3 and HA PEDV N constructs were co transfected into HEK293T and LLC PK1 cells, respectively. At 24 h post transfection, protein samples were harvested for Co IP assays. Given that both proteins are RNA binding proteins, 1 µL of RNase A was added and incubated for 30 min prior to elution. Western blotting analysis after elution revealed their interaction. (C, D) Flag RBM3 and HA PEDV N constructs were co transfected into HeLa and LLC PK1 cells, respectively. At 24 h post transfection, confocal microscopy was performed. Fluorescent signal acquisition was conducted on a confocal laser scanning microscope following DAPI nuclear counterstaining. The scale bar represents 50 μm.(E) In LLC PK1 cells overexpressing Flag RBM3, cells were infected with PEDV and harvested at 4 h post infection for confocal microscopy. Fluorescent signal acquisition was conducted on a confocal laser scanning microscope following DAPI nuclear counterstaining. The scale bar represents 50 μ m.(F) Flag RBM3 and HA NSP12 constructs were co transfected into LLC PK1 cells for Co IP assays. Prior to elution, 1 µL of RNase was added and incubated for 30 min. Western blotting analysis after elution showed no interaction.

### 3.4 RBM3 increases HSPA5 expression to promote PEDV proliferation

Given that RBM3, as an RNA-binding protein, not only acts directly on viral RNA but may also regulate host gene expression, and that HSPA5 has been reported as a key positive host factor for PEDV entry [12], we further explored the relationship between RBM3 and the host protein HSPA5. We either overexpressed or knocked down RBM3 in LLC-PK1 cells, and RNA was extracted 24 h later. RT qPCR analysis revealed that RBM3 perturbation altered HSPA5 mRNA levels (Fig. 4A, B), suggesting that RBM3 may regulate HSPA5 transcription. To further investigate this, we cloned truncated fragments of the HSPA5 promoter into the pGL3 basic vector (Fig. S2A) and examined their promoter activity using a dual luciferase reporter assay (Fig. S2B). HEK293T and LLC-PK1 cells were separately co-transfected with the validated promoter construct, the Flag-RBM3 plasmid, and the Renilla luciferase internal control plasmid. In both cell lines, the assay showed that RBM3 overexpression significantly increased HSPA5 promoter activity compared with the control group (Fig. 4C, D). Given the role of RBM3 in stabilizing viral RNA, to rule out the possibility that RBM3-mediated upregulation of HSPA5 is related to this function, we designed the following experiments. First, cell viability was assessed by CCK-8 assay (Fig. S2C) and the amount of actinomycin D (2.4 μM) was determined. Next, we overexpressed RBM3 in LLC-PK1 cells, while control cells were transfected with the empty vector pXJ41. At 24 h post-transfection, both the experimental and control cells were treated with actinomycin D. RNA samples were collected starting at 4 h after treatment and then every 2 h. RT-qPCR was used to measure endogenous HSPA5 mRNA levels, and the decay curve was generated accordingly. Quantification of the decay curves showed that the degradation rate constant () did not differ significantly between control cells and cells overexpressing RBM3 (Fig. S2D). These data make it unlikely that RBM3 stabilizes HSPA5 mRNA.

**Figure 4.**
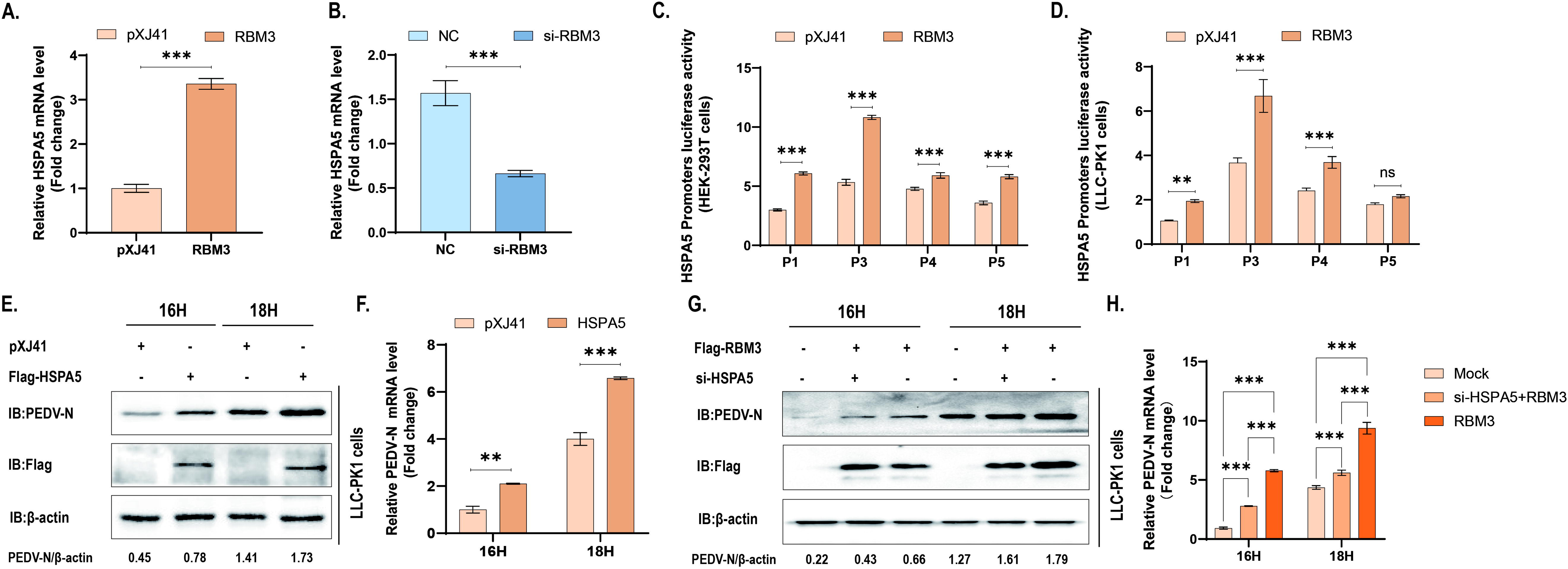
RBM3 increases HSPA5 expression to promote PEDV proliferation. (A, B) RT qPCR analysis of endogenous HSPA5 mRNA levels in LLC PK1 cells following RBM3 overexpression or knockdown, respectively. P-values determined by two-way ANOVA. (C, D) The truncated HSPA5 promoter constructs and the internal control plasmid were co transfected into HEK293T and LLC PK1 cells, respectively, with Flag RBM3 overexpressed in the experimental group and empty vector in the control group. Dual luciferase reporter assays were performed to evaluate the effect of RBM3 overexpression on HSPA5 promoter activity. P-values determined by two-way ANOVA. (E, F) Western blotting and RT qPCR analysis of cells overexpressing Flag HSPA5 versus empty vector control at indicated hours post infection (hpi) with PEDV. P-values determined by two-way ANOVA. (G, H) LLC PK1 cells were transfected with si HSPA5 plus Flag RBM3, or NC plus Flag RBM3, then infected with PEDV. Western blotting and RT qPCR analyses were performed at indicated time points compared to the vector control. P-values determined by two-way ANOVA. All data represent three independent biological replicates (n=3). Error bars represent mean ± SD. *p < 0.05, **p < 0.01, ***p < 0.001.

To confirm the proviral function of HSPA5 in our experimental system, we repeated the phenotypic experiments of HSPA5 and PEDV in LLC PK1 cells. Both Western blotting (Fig. 4E) and RT qPCR (Fig. 4F) results confirmed that HSPA5 promotes viral proliferation. To this end, we sought to define the role of HSPA5 in RBM3-mediated promotion of viral proliferation, LLC-PK1 cells were co-transfected with pXJ41+NC (mock control), si-HSPA5+Flag-RBM3, or NC+Flag-RBM3. At 24 h post-transfection, the cells were infected with PEDV, and samples were collected at the indicated time points for Western blotting and RT-qPCR. The results showed that RBM3 overexpression still promoted viral proliferation in the presence of HSPA5 knockdown, although the effect was attenuated compared with that in the absence of HSPA5 knockdown (Fig. 4G, H). The knockdown efficiency of si-HSPA5 was confirmed by RT-qPCR (Fig. S2E).

## 4. Discussion

Host RNA-binding proteins (RBPs) have emerged as key regulators of the PEDV life cycle. For instance, LARP4 cooperates with PABPC1 to bind the viral 3’UTR and promote translation efficiency [13], while DDX5 and SND1 facilitate RTC assembly and viral RNA synthesis through interactions with both viral RNAs and non-structural proteins [3]. RBM3 is a cold-shock protein with reported RNA chaperone-like activity and has been shown to enhance the stability of multiple host mRNAs, such as MEF2C [14], but its role in coronavirus infection has not been explored. In this study, we identified a dual proviral role for RBM3 in PEDV infection. At the direct level, RBM3 preferentially bound to the PEDV 3’UTR, which was associated with prolonged viral RNA half-life and increased viral RNA accumulation. In addition, RBM3 interacted with the PEDV N protein and dsRNA, but not with NSP12, a core component of the replication-transcription complex (RTC). Whether these interactions functionally contribute to RTC activity or viral RNA synthesis remains to be further investigated. At the indirect level, RBM3 increases the expression of HSPA5, a host factor known to facilitate PEDV attachment and internalization by binding to the spike protein [12]. These two mechanisms likely act at distinct stages of the viral life cycle and collectively contribute to efficient PEDV proliferation.

RBM3 is well characterized as an RNA chaperone that maintains host mRNA stability and translation efficiency under stress conditions such as hypothermia and hypoxia [8]. In the context of PEDV infection, our data revealed that RBM3 associates with PEDV RNA, with a preferential binding to the 3’UTR region (Fig. 2A-C), which is known to be closely linked to RNA stability [15]. To further investigate the functional significance of this association, we performed functional assays, which demonstrated that RBM3 overexpression extended the viral RNA half life from 4.14 h, 4.21 h, 4.936 h to 7.75 h, 7.17 h (Fig. 2D) and increased the initial accumulation of viral RNA during the early stage of viral proliferation (Fig. 2E, F). These findings are consistent with the canonical function of RBM3 as an RNA stabilizing chaperone, while extending its substrate repertoire from host mRNAs to viral RNAs. Interestingly, we also found that RBM3 interacts with the PEDV N protein and colocalizes with dsRNA (Fig. 3A-E), suggesting its potential involvement in the processing of viral replication intermediates. Notably, however, RBM3 failed to interact with NSP12 (Fig. 3F), the core component responsible for PEDV RNA synthesis [16]. Based on these findings, we propose a speculative working model in which RBM3 may be recruited to the replication transcription complex (RTC) through its interaction with the N protein, where it exerts its RNA chaperone activity in situ to potentially protect viral RNA from degradation and facilitate processing of replication intermediates. The lack of direct interaction between RBM3 and NSP12 is consistent with the possibility that RBM3 does not directly participate in the catalytic process of RNA synthesis, but may instead serve as an “auxiliary factor” or “structural scaffold” for the RTC. In this capacity, RBM3 could potentially enhance viral RNA synthesis by stabilizing viral RNA substrates or facilitating their transition from the replication complex to downstream translation or packaging. Nevertheless, this model remains highly speculative and awaits further experimental validation.

RBM3 is also known to play roles in cellular stress responses, such as modulating PERK phosphorylation via NF90 [17] and promoting AKT activation under cold exposure [18], and whether RBM3 influences viral infection through regulation of other host proteins has remained unexplored. One such host protein is HSPA5 (GRP78/BiP), a well-established endoplasmic reticulum chaperone [19] that can translocate to the cell surface and serve as an attachment or entry factor for multiple coronaviruses, including SARS-CoV-2, MERS-CoV, and PEDV [12, 18, 19]. In PEDV infection, HSPA5 binds directly to the spike protein and promotes viral internalization [12]. In this study, we found that RBM3 increases HSPA5 expression, thereby indirectly promoting PEDV infection. RT-qPCR analysis revealed that modulation of RBM3 altered HSPA5 mRNA levels (Fig. 4A, B), and dual-luciferase reporter assays showed that RBM3 overexpression increased HSPA5 promoter activity (Fig. 4C, D), suggesting that RBM3 may regulate HSPA5 at least in part at the transcriptional level. To further dissect the regulatory axis between these two proteins, we performed a series of mechanistic investigations. First, given the known RNA stabilizing function of RBM3, we employed the transcriptional inhibitor actinomycin D and constructed an HSPA5 mRNA decay curve (Fig. S2D), which ruled out the possibility that RBM3 upregulates HSPA5 through enhanced mRNA stability. Second, we co transfected LLC PK1 cells with si HSPA5 + Flag RBM3 or NC + Flag RBM3, with pXJ41 + NC serving as a mock control, and analyzed the corresponding phenotypes. Compared with the mock control, the si HSPA5 + RBM3 group still exhibited significant viral proliferation, whereas the group without HSPA5 knockdown showed an even more pronounced enhancement of viral proliferation (Fig. 4G, H). These results indicate that even after HSPA5 knockdown, RBM3 overexpression retained partial proviral activity, suggesting that the direct RNA stabilizing effect operates independently of HSPA5. Conversely, the attenuated proviral effect observed upon HSPA5 depletion confirms that HSPA5 upregulation contributes significantly to the overall proviral function of RBM3. The RBM3 HSPA5 regulatory axis has significant biological implications during PEDV infection. From the host perspective, RBM3 mediated regulation of HSPA5 could represent a protective stress response aimed at restoring ER homeostasis and limiting virus induced cellular damage. Paradoxically, however, this protective response is exploited by the virus: the increased surface availability of HSPA5 resulting from RBM3 mediated upregulation predisposes cells to PEDV infection, potentially lowering the threshold for spike protein dependent viral entry. Importantly, our results place RBM3 upstream of HSPA5 for the first time, revealing a functional link between these two proteins. This finding expands the known functional spectrum of RBM3 (from direct RNA metabolism to the regulation of host entry factors) and provides a novel perspective on how a single host RBP can modulate viral infection through both direct and indirect mechanisms. However, the lack of HSPA5 Western blotting data means protein-level changes remain unconfirmed, representing a limitation.

In summary, this study reveals a dual regulatory mechanism of RBM3 in PEDV replication: a direct pathway that stabilizes viral RNA, thereby increasing the accumulation of viral RNA, and an indirect pathway in which RBM3 increases HSPA5 expression, thereby rendering cells more susceptible to PEDV infection. These two pathways likely operate at distinct stages of the viral life cycle— entry and replication— and collectively contribute to efficient PEDV proliferation. Our findings have several implications. First, they identify RBM3 as a positive host regulator of the alphacoronavirus PEDV, expanding our understanding of cold-shock protein functions in enteric coronavirus infection. Second, the newly established functional link between RBM3 and HSPA5 suggests the existence of a regulatory axis that may offer a theoretical basis for developing antiviral strategies targeting both viral entry and replication. Additionally, given the conserved nature of the 3’UTR among coronaviruses, the ability of RBM3 to recognize and stabilize the PEDV 3’UTR raises the possibility that RBM3 may act as a potential broad-spectrum regulator of coronavirus RNA stability, a hypothesis that warrants future investigation in other coronaviruses such as SARS-CoV-2. Collectively, these results provide new molecular insights into virus-host interactions and may offer clues for the future design of anti-coronavirus therapeutic strategies.

## Declarations

### Ethics approval and consent to participate

Not applicable.

### Consent for publication

Not applicable.

### Availability of data and materials

The datasets used and analysed during the current study are available from the corresponding author on reasonable request.

### Authors’contributions

Conceptualization: S.L., Y.L.; Methodology: S.L., Y.L., X.Y., N.K.; Investigation: S.L., J.L., W.Q., Y.Z., H.L.; Writing-original draft: S.L., N.K., Y.Z.; Writing-review & editing: S.L., N.K., X.C., G.T., T.L., Z.C. J.L., and Y.Z.; Supervision: T.S.; All authors reviewed the final manuscript.

## Supporting information

Supplementary data for Figure 2

Supplementary data for Figure 4

## Acknowledgements

Not applicable.

---

**Figure S1. Supplementary data for Figure 2.**

(A) Cells were transfected with plasmids harboring the PEDV 3’UTR fragment or the N plasmid, respectively, and the impact of RBM3 overexpression or knockdown on the corresponding RNA abundance was examined by RT qPCR. P-values determined by two-way ANOVA. (B) CCK 8 assay was performed to evaluate cell viability upon treatment with varying doses of Remdesivir, indicating that cell viability remained above 90% at both 10 µM and 20 µM.(C) LLC PK1 cells were infected with PEDV and subsequently treated with 20 µM Remdesivir, viral RNA levels were measured at different time points post treatment to determine the time point (T0) at which RNA synthesis had completely ceased. RT-qPCR result revealed that T0 fell between 6 and 9 h. All data represent three independent biological replicates (n=3). Error bars represent mean ± SD. *p < 0.05, **p < 0.01, ***p < 0.001.

**Figure S2. Supplementary data for Figure 4.**

(A) Schematic diagram of truncated HSPA5 promoter fragments cloned into the pGL3 basic vector.(B) Dual luciferase reporter assay to assess the promoter activity of each truncated fragment, confirming the construct with verified activity.(C) CCK 8 assay to evaluate cell viability upon treatment with varying doses of actinomycin D.(D) In Flag RBM3 overexpressing cells and vector control cells, actinomycin D (2.4 µM) was added, and samples were collected every two hours starting from 4 h post treatment. RT qPCR was performed to analyze endogenous HSPA5 mRNA levels and generate the decay curve. The did not differ significantly between the control group and the experimental group. P-values determined by unpaired t-test. P=0.058. (E) Assessment of the interference efficiency of si-HSPA5. P-values determined by unpaired t-test. All data represent three independent biological replicates (n=3). Error bars represent mean ± SD. *p < 0.05, **p < 0.01, ***p < 0.001.

