## Supplementary figures and images for "Dual role of RBM3 in PEDV infection: stabilization of viral RNA and upregulation of the entry factor HSPA5"

### Figure.S1.jpg

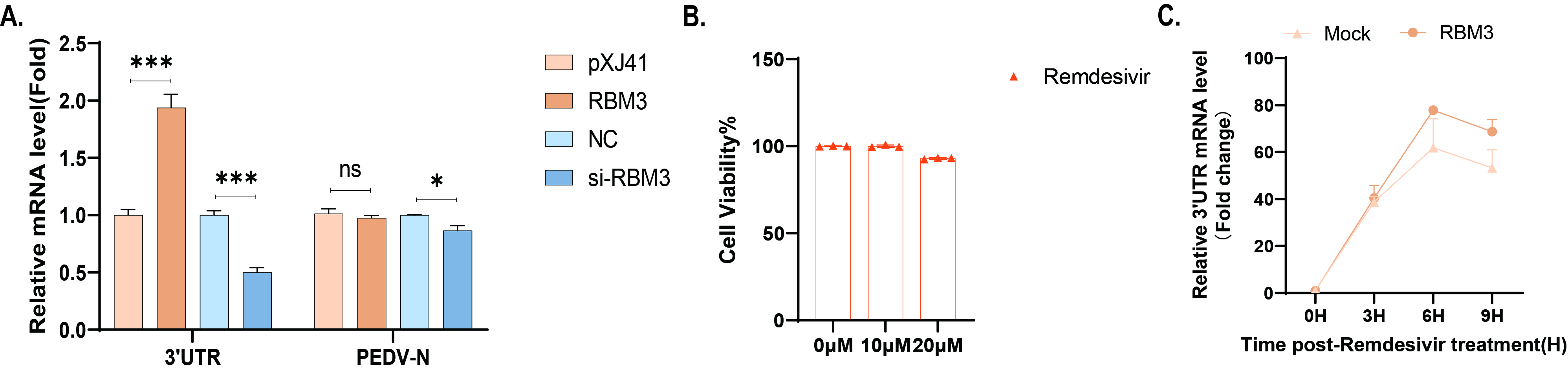

### Figure.S2.jpg

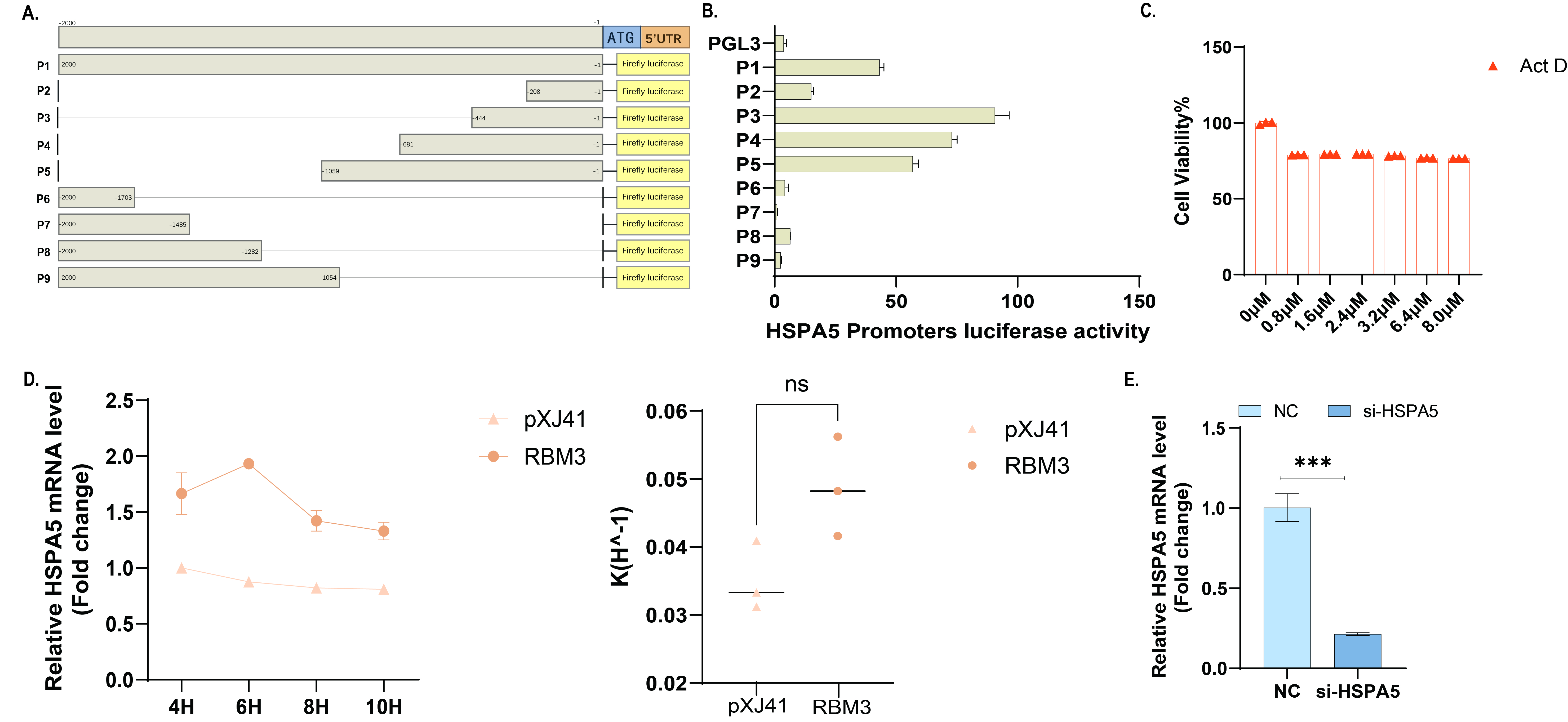
